# Pathway-guided deep neural network toward interpretable and predictive modeling of drug sensitivity

**DOI:** 10.1101/2020.02.06.930503

**Authors:** Lei Deng, Yideng Cai, Wenhao Zhang, Wenyi Yang, Bo Gao, Hui Liu

## Abstract

**Motivation:** To efficiently save cost and reduce risk in drug research and development, there is a pressing demand to develop *in-silico* methods to predict drug sensitivity to cancer cells. With the exponentially increasing number of multi-omics data derived from high-throughput techniques, machine learning-based methods have been applied to the prediction of drug sensitivities. However, these methods have drawbacks either in the interpretability of mechanism of drug action or limited performance in modeling drug sensitivity.

**Results:** In this paper, we presented a pathway-guided deep neural network model, referred to as pathDNN, to predict the drug sensitivity to cancer cells. Biological pathways describe a group of molecules in a cell that collaborates to control various biological functions like cell proliferation and death, thereby abnormal function of pathways can result in disease. To make advantage of both the excellent predictive ability of deep neural network and the biological knowledge of pathways, we reshape the canonical DNN structure by incorporating a layer of pathway nodes and their connections to input gene nodes, which makes the DNN model more interpretable and predictive compared to canonical DNN. We have conducted extensive performance evaluations on multiple independent drug sensitivity data sets, and demonstrate that pathDNN significantly outperformed canonical DNN model and seven other classical regression models. Most importantly, we observed remarkable activity decreases of disease-related pathway nodes during forward propagation upon inputs of drug targets, which implicitly corresponds to the inhibition effect of disease-related pathways induced by drug treatment on cancer cells. Our empirical experiments show that pathDNN achieves pharmacological interpretability and predictive ability in modeling drug sensitivity to cancer cells.

**Availability:** The web server, as well as the processed data sets and source codes for reproducing our work, is available at http://pathdnn.denglab.org

## 1 Introduction

The incidence of cancer increases gradually as people getting older, posing a serious threat to human health [1, 2, 3], thereby leading to a growing demand for anticancer drugs [4]. Unfortunately, traditional anti-cancer drug research and development, such as active ingredient identification, is a high-cost and high-risk task [5]. Therefore, it is urgent to develop systematic methods that can integrate and exploit the chemical, pharmacological and genomic data resource to accurately predict drug sensitivity to cancer cells.

In the past few years, *in silico* drug discovery has progressively evolved and many computational methods that are more targeted and efficacious have been proposed [6, 7]. These methods are mainly based on molecule docking, data mining and machine learning techniques, among which machine learning-based methods have drawn intensive attention from research community and gained encouraging performance. For example, Menden *et al.* proposed a machine learning method to predict the interactions between drugs and cancer cells based on genomic and chemical properties [8]. It was designed to optimize the design of drug and cell line screening experiments by detecting thousands of drugs through systematic test of their potential as anti-tumor therapies. Neto *et al.* presented a Bayesian inference method, referred to as STREAM [9], which averaged on the adjusted parameter grid for efficient parameter optimization and adopted singular value decomposition reparameterization of ridge regression model for high efficient matrix inversion operation. However, a large number of features were excluded during feature extraction process so that its prediction reliability is not convincing enough. Raziur *et al.* subsequently proposed another method based on random forest to predict drug sensitivity, in which the regression tree is expressed in the form of a probability tree and the nature of heteroscedasticity is analyzed [10]. As a result, the prediction set synthesizes both mixed distribution and weighted sum of relevant random variables, which is stated to produce reliable prediction. Nonetheless, during the re-implementation of this method, we found it is time-and-memory resource consuming, which makes its scalability limited.

In fact, the drug sensitivity is determined by multiple factors, such as binding affinity between drug molecules and targets, mechanism of drug action and resistance of cancer cells to drug administration [11, 12]. A feasible method to improve the prediction of drug sensitivity is to comprehensively take into account many biomarkers [13]. Nonetheless, the features when using multiple biomarkers are huge and complicated, which makes it difficult to be applied by traditional machine learning methods that performed well on small datasets and limited features [14]. For the capacity to learn higher level of feature representation, deep learning has become an effective and popular tool in many fields, including Bioinformatics community [15]. For example, Wang et al. proposed a pairwise input neural network for target-ligand interaction prediction [16]; Aliper *et al.* used deep learning to predict pharmacological properties of drugs based on transcriptomic data [14]; Wan et al. also successfully predicted the compound-protein interactions by deep learning [17].

Despite high accuracy and compatibility of high-dimensional data, existing deep learning-based methods for drug sensitivity function as “black boxes”, lack of interpretability in terms of biological and pharmacological views. In this paper, we presented a pathway-guided deep neural network model, referred to as pathDNN, to predict the drug sensitivity to cancer cells. As drug molecules interact with their targets and subsequently inhibit or activate targeted pathways, which eventually lead to phenotypical change of the cancer cells. To explore the pharmacological mechanism of action during the drug treatments, we take the known biological pathways into account, as well as genome-wide expression profiles of cancer cell lines, drug-target interactions and large-scale quantitative drug sensitivity datasets, to construct a unified computational framework for predicting drug sensitivity. Specifically, we reshape the canonical DNN structure by introducing a layer of pathway nodes and their connections to input gene nodes and drug target nodes. The pathway layer is followed by several fully-connected layers and an output layer. We conducted extensive performance evaluations on multiple independent drug sensitivity data sets, and demonstrate that pathDNN significantly outperformed not only canonical DNN model, but also seven other classical regression models. Most importantly, we observed remarkable output decreases of disease-related pathway nodes during forwarding propagation upon inputs of drug targets, which implicitly corresponds to the inhibition effect of disease-related pathways induced by drug treatment on cancer cells. Our empirical experiments show that pathDNN successfully makes advantage of both the excellent predictive ability of deep neural network and the biological knowledge of pathways, and achieves pharmacological interpretability and predictive ability in modeling drug sensitivity to cancer cells.

## 2 Methods

### 2.1 Data Resource

#### Drug Sensitivity

The drug sensitivity data were obtained from GDSC [18], which assayed 250 anti-cancer drugs on 970 cancer cell lines. In total, the GDSC drug sensitivity dataset includes 198,929 drug response data points, from which the area under the drug response curve (Actarea value) is computed as the quantitative response metric to measure the sensitivities of drugs to cancer cell lines. Intuitively, the higher the Actarea value, the more sensitive the drug to cancer cell line, namely, the higher the capability of the drug to kill cancer cells. We normalized the Actarea value with the Min-Max scaling method so that the Actarea values fall into the range [0,1].

#### Pathway maps

The pathway maps were obtained from Kyoto Encyclopedia of Genes and Genomes (KEGG) [19], a public database devoted to the understanding of high-level functions and utilities of the biological system. Each KEGG pathway is a collection of the manually drawn maps representing current knowledge on the molecular interaction, reaction and relation networks, covering metabolism, cellular processes, organismal systems and human diseases. The current repository contains 323 common signaling pathways involved in 7,230 unique genes, each pathway map involves tens of genes, and each gene may be engaged in one or more pathways.

#### Drug targets

We downloaded drug-protein interactions from STITCH database [20], which is a comprehensive database that collected compound-protein interactions from different sources: biochemical experiments, external databases, text mining and computational predictions. Meanwhile, STITCH has computed a confidence score for each interaction ranging from 0 to 1,000, which indicates the confidence of the compound-protein interaction supported by four types of evidence. Because we supposed that too low-confidence targets are probable unauthentic ones, we used a confidence threshold of 0.8 (corresponding to 800 combined score in STITCH) to remove low-confidence target proteins, In total, 250 drugs and 1100 target proteins are derived from the STITCH dataset.

#### Expression profiles

To characterize the cancer cell lines, we downloaded the expression profiles of cancer cell lines from GDSC, which has profiled the expression levels of the cancer cell lines used in the drug sensitivity assays. The expression profiles cover 17,737 genes on 970 cancer cell lines. The expression levels have been adjusted by Robust Multi-array Average (RMA) method to remove the background noise. Subsequently, we centered the expression levels over genes for each cancer cell line so that the mean is zero.

### 2.2 Data preprocessing

To reduce the dimensionality of input features, the set of “landmark genes” was adopted in our data preprocessing for feature selection. According to the conclusion of LINCS Project, approximately 80% of the information can be captured by 977 landmark genes, thereby we employed these genes in our analysis rather than the whole genome-wide genes.

For the 250 drugs assayed in the GDSC drug sensitivity dataset, 1100 proteins targeted by these drugs are obtained, and the protein identifiers were converted to gene identifiers consistent with KEGG pathway identifiers. Subsequently, the set of genes were intersected with those included in the pathway dataset and landmark genes, and 741 genes were filtered out and used as input perturbation on the molecule network. Similarly, 17,737 genes covered in the genome-wide expression profiles of cancer cell lines were intersected with those in the pathway and landmark gene dataset. Hence, 537 genes were extracted to characterize the genotype of cell lines from the perspective of expression profiles. Finally, two types of gene sets, representative features of drug perturbation and genotypes of cell lines, were concatenated to form the input layer of 1,278 nodes, with which our model integrates both drug targets and expression profiles of cancer cells. Figure 1 shows the numbers of each data set and overlaps among them, the final overlapping genes concatenates to act as the input layer composed of drug targets and expression profiles.

**Fig. 1:**
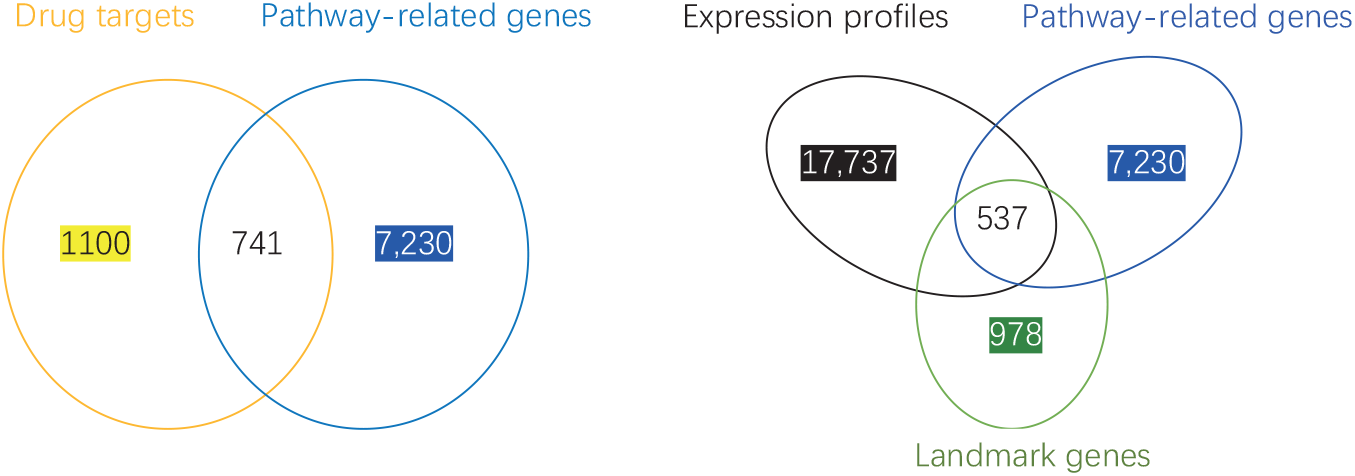
Number of genes in drug targets, pathways and gene expression profiles and overlaps among them.

### 2.3 Pathway-guided deep neural network

Signaling pathways are functional modules that control the cellular processes ranging from cell proliferation and survival to specialized functions such as the immune response and angiogenesis. Pathway maps are the representations of our understanding of various biological processes. In drug treatment on cancer cells, drug molecules interact with their targets and subsequently inhibit or activate relevant pathways, which eventually lead to a phenotypical change of the cancer cells. To explore the pharmacological mechanism of action, we take the pathways to construct a DNN model, by introducing a layer of pathway nodes and their connections to input gene nodes. The pathway layer is followed by two fully-connected hidden layers and an output layer. As shown in Figure 2, each neuron in the input layer represents a gene (cell line feature or drug target), while the corresponding output is the quantitative drug sensitivity. In particular, for a drug tested on a cell line, the input values of cell line nodes are normalized gene expression levels, and the input values of drug target nodes are the STITCH confidence scores if targeted and zero if not targeted by the assayed drug. The first hidden layer, referred to as the pathway layer, has 323 neurons derived from the KEGG pathway repository. The connections between the input layer and the pathway layer are determined by the associations between genes and pathways. A 323*1278 mask matrix, denoted by *M*, was used to encode the relation between gene nodes and pathway nodes, in which 1 represents the existence of an association between the gene and pathway node, and 0 otherwise. In the back propagation process, the weights of the edges are iteratively updated using the rule as below:

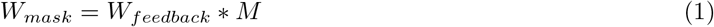

where *W*_*feedback*_ represents the feedback weights in last backpropagation iteration, *W*_*mask*_ represents the successive forward weight which asserts the sparsity of gene-pathway association. Particularly, in the first iteration, *W*_*feedback*_ is the originally assigned weight. A sequence of fully-connected hidden layers is successively presented. Let *a*_*n*_ be the vector of hidden layer *H*_*n*_, the feed-forward process can be iteratively calculated as below:

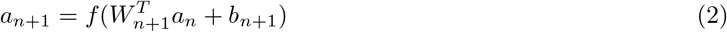

where *W*_*n*+1_ is the weights between hidden layer *H*_*n*_ and *H*_*n*+1_, *b*_*n*+1_ is the bias vector, *f* is the activation function that performs nonlinear transformations. To avoid vanishing gradient problem [21], the rectified linear unit (ReLU) was used as the activation function [22]. Mean square error (MSE) that measures the average of the squares of errors, was applied as the loss function to indicate the degree of convergence. The MSE is defined as follows:

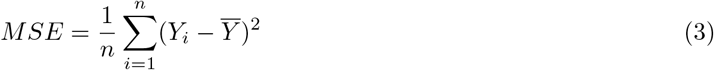

in which *Y*_*i*_ represents the true values and *Y̅* represents the predicted outputs. By minimizing the loss function through optimization algorithms, our pathDNN model can be iteratively optimized by updating weights and biases.

**Fig. 2:**
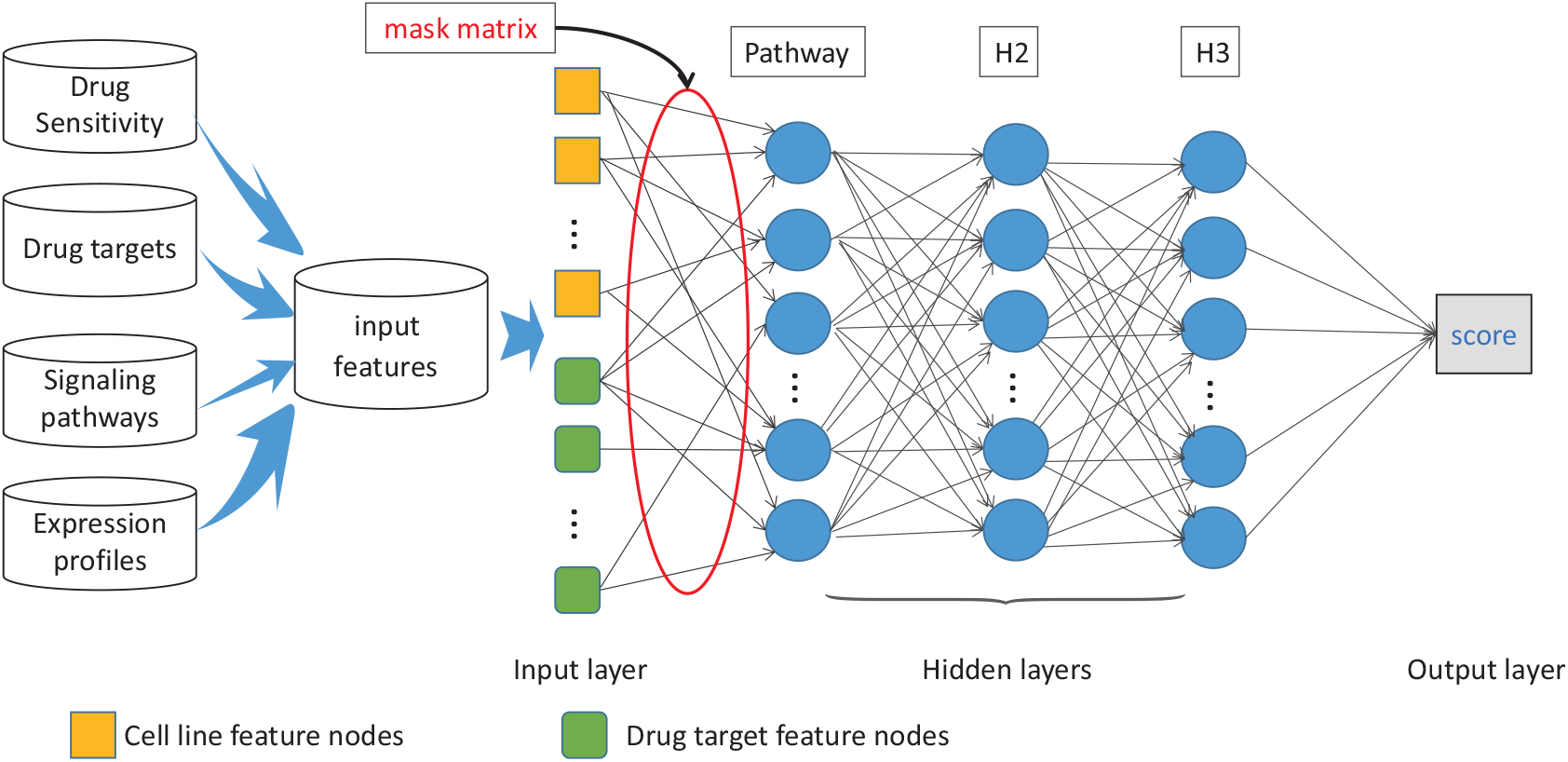
Illustrative diagram of the data sources and structure of the pathway-guided deep neural network. The first hidden layer is composed of the pathway nodes that connected to the input genes compiled from genome-wide expression profiles and drug targets, the mask matrix is used to encode the relation between gene nodes and pathway nodes. The second and third hidden layers are fully-connected, and single node in the output layer output the quantitative drug sensitivity.

## 3 Results

### 3.1 Performance measurement

Since MSE loss reflects the goodness of fit and the curve of MSE can indicate the training speed, it was chosen as one performance measure to evaluate our proposed method. The 10-fold cross-validations were carried out to evaluate the overall performance. The training set is randomly split into ten subsets of roughly equal size. Each subset, in turn, is taken as the test set, and then a predictive model is trained on the samples included in the rest nine subsets, and is evaluated on the test set. As a result, the Pearson correlation coefficient (PCC) between the predicted and real drug sensitivities was calculated to evaluate the performance. The PCC is formulated as below:

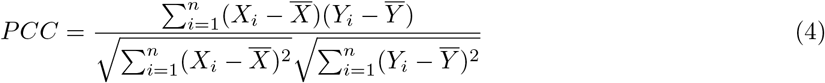

where (*X_i_*, *Y*_*i*_) represents a set of predicted and true drug sensitivity, *X̅* represents the mean of the predicted sensitivity, *Y̅* representing the mean of the true sensitivity. PCC∈[−1, 1], the closer to 1 of the absolute PCC value, the higher the correlation between predicted and true sensitivity, and 0 otherwise.

### 3.2 Hyperparameter tuning

All of our experiments were implemented on a CentOS Linux server equipped with 4 NVIDIA Corporation GM107GL graphics cards that have 32G video memory in total. Pytorch, an open-source machine learning library for Python and one of the most prevalent deep learning frameworks, was the main DNN dependency in our work. Thanks to its intuition and flexibility, we could conduct the experiments with high efficiency. GPU acceleration technique, which can handle the tensors at an incredible level, was also used to accelerate the training speed.

To tune the hyperparameters, we first roughly adjusted the hyperparameters by grid search and obtained their candidate values or intervals, and then fine tuned the parameters around the rough values or in the intervals to achieve the best performance. The self-adapted optimizer Adam [23] was firstly used to pre-train the model and then stochastic gradient descent (SGD) [24] was employed for further training. Since the SGD optimizer will not be applied used until the model mainly converges, the learning rate of SGD is fixed at a relatively tiny value 0.01, and the weight decay is set as 0.001. Therefore, the hyperparameters to be tuned include the number of hidden layers, the size of each hidden layer, the learning rate and the weight decay of Adam optimizer, the batch size and the epoch number. We implemented 10-fold cross-validation on the benchmark set, with the PCC value to evaluate the performance during hyperparameter tuning.

As shown in Figure 3, the number of batch size mainly affect the training time, while other parameters significantly influence the performance. We empirically set these parameters to the values achieving the best performance. As a result, the learning rate of Adam optimizer is set to 0.0001, the batch size is set to 512, the weight decay and epoch of Adam optimizer are set to 0.0001 and 500, respectively.

**Fig. 3:**
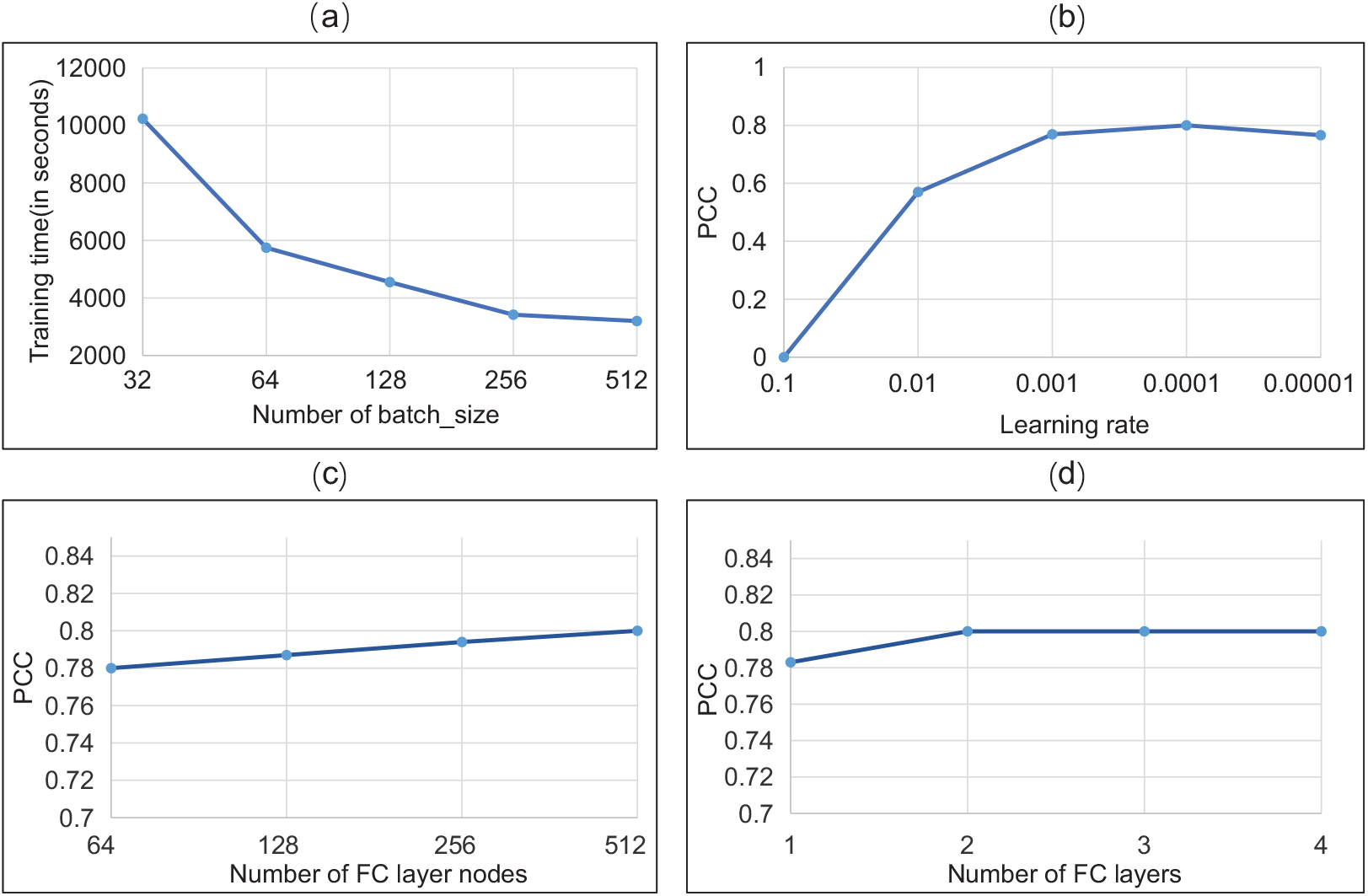
Impact of four hyperparameters on the performance of pathDNN model

### 3.3 Comparison with competitive methods

We subsequently carried out performance evaluation by comparing pathDNN with both traditional machinelearning classifiers, including classical linear regression, lasso regression and random forest, and a couple of existing methods, including scalable-time ridge estimator by averaging of models (STREAM) [9], efficient neural network (Enet) [25], weighted random forests (WRF) [26] and probabilistic random forests (PRF) [10]. As shown in Fig 4, pathDNN is far superior to all competitive methods in terms of prediction accuracy. STREAM uses SVD reparameterization to obtain the improvement of computational efficiency. However, a large number of features were excluded during the feature selection so that its prediction performance is not satisfactory. As to WRF and PRF, these two methods are time-consuming on the operation. Therefore, our proposed pathDNN is believed to be superior to all aforementioned methods, with respect to both time and prediction accuracy.

**Fig. 4:**
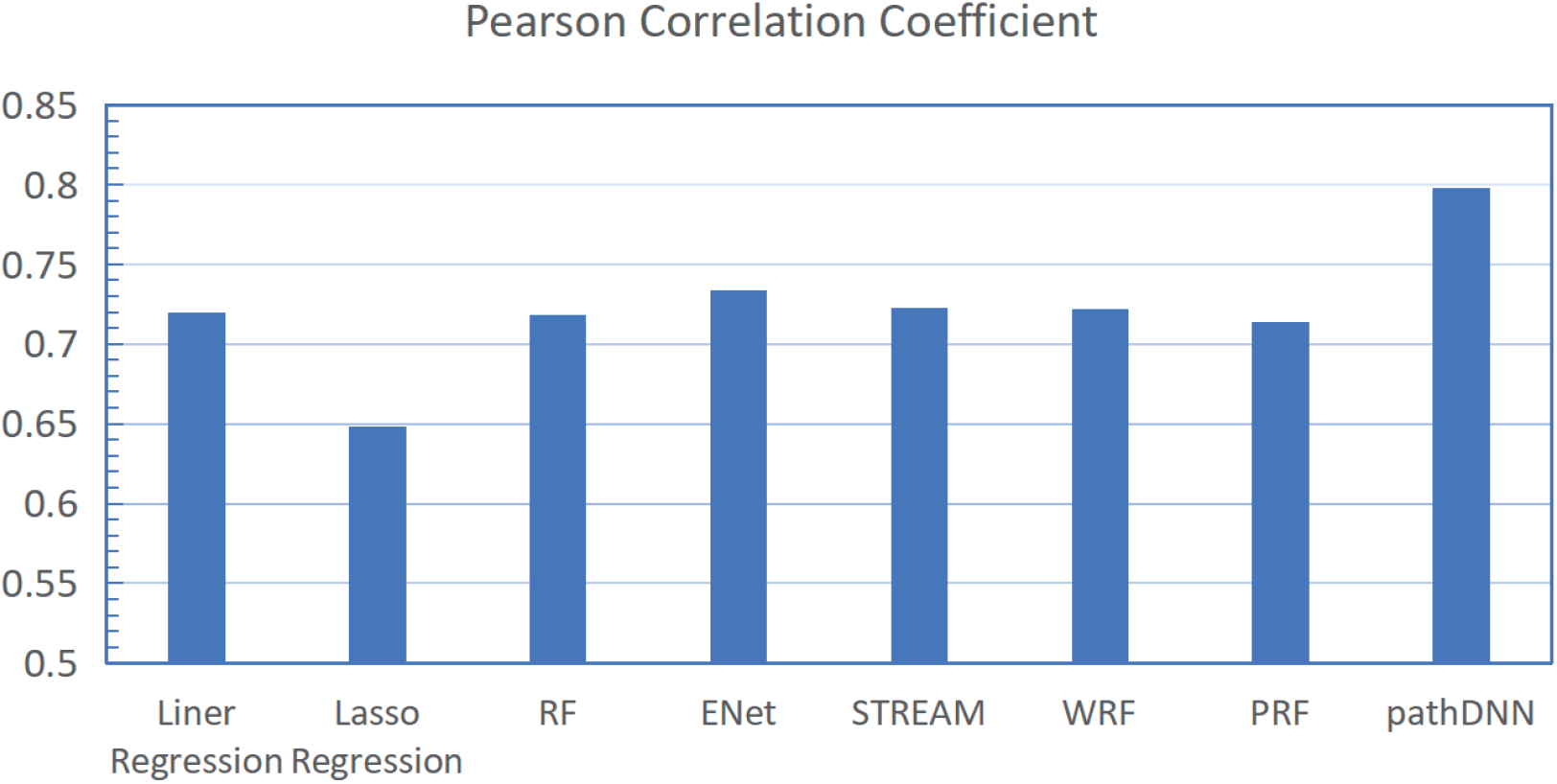
Performance comparison to seven existing methods on GDSC drug sensitivity dataset.

### 3.4 Leave one cell line out validation

To explore the predictive power in finding new indications of approved drugs, we performed leave-one-out validation by iteratively removing one cancer cell line from training set and then test it on the trained model, which is referred to as leave one cell line out validation. Note that all drug sensitivities in association to the leave-out cell line are excluded from the training set, and then the Pearson correlation coefficients between the predicted and real sensitivities regarding each cell line was computed. To illustrate the experimental results, we presented a scatter plot to exhibit the results performed by leave-one-cell-line-out validation. As demonstrated in Figure 5, our model obtained notably high correlation coefficient on most cell lines. Specifically, PCC≤0.5 accounted for 1.24%, 0.5≤PCC*<*0.7 accounted for 20.93, 0.7≤PCC*<*0.8 accounted for 38.56%, and PCC≥0.8 accounted for 39.28%.

**Fig. 5:**
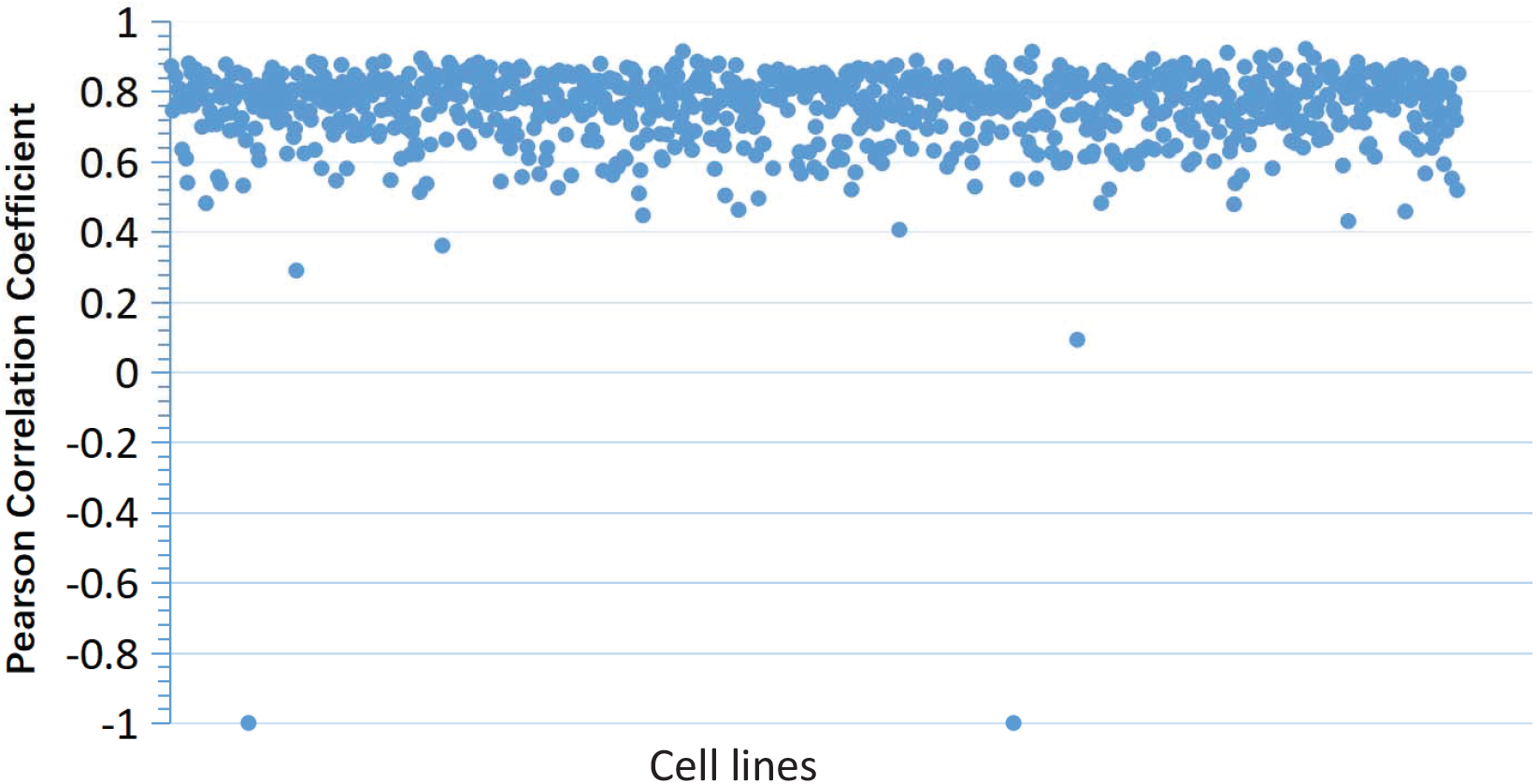
Performance evaluation of pathDNN model by leave one cell line out validation.

### 3.5 Evaluation on independent dataset

To further evaluate the performance of pathDNN, we employed another drug sensitivity data set released by the Cancer Cell Line Encyclopedia (CCLE) project [27], which assayed the response levels of 491 cancer cell lines to 22 small molecule drugs. Together with drug-target interactions and pathways mentioned above, we constructed another independent dataset for performance evaluation. The mask matrix encoding the connections between the input nodes and pathway layer nodes is the same as the one mentioned above. On this independent dataset, we conducted a 10-fold cross-validation experiment and our model achieved a PCC value of 0.91.

Also, we use genomic variants derived from high-throughput sequencing of cancer cell lines instead of gene expression profiles. For genomic variants, we set the input values of genes with one or more variants as 1, and 0 otherwise. We performed 10-fold cross-validation based on the variants dataset and obtained 0.013 MSE value and 0.81 PCC value, implying that pathDNN can still achieve high performance. Meanwhile, our empirical experiments showed that expression profiles are more informative than genomic variants in predicting drug sensitivity to cancer cells.

To further evaluate the superiority of our model, we trained the DNN model based on the CCLE dataset and then tested on the GDSC dataset. In order to ensure that the genes selected as input features of the two data sets are consistent, we took the intersection of the characteristic genes as the input nodes. This experiment cross different datasets showed 0.05 MSE value and 0.75 PCC value, which effectively demonstrated the excellent generalization capacity of our model.

### 3.6 Pathway layer promoting performance and interpretability

To test whether the pathway layer contributes to the improvement of performance or not, we took a comparison to the classical fully-connected DNN model of which the hyperparameters have been optimized by grid search and model parameters have been tuned to reach the best goodness of fit. The results demonstrated that pathDNN significantly outperformed classical fully-connected DNN model, i.e., pathDNN reaches 0.80 PCC value on 10-fold cross validation, whereas the fine-tuned canonical DNN model can only obtain 0.73 PCC value.

To explore the interpretability of our model, we compared the outputs of the pathway nodes with drug treatments to those without drug treatments (corresponding drug target feature nodes set to 0). Expectedly, we observed remarkable decreases in disease-related pathway nodes upon drug treatments. Taking the disease rhabdomyosarcoma as an example, drug CTK7H7014 (CHEMBL1623573) is approved for its blocking of the hsa05202 pathway (the transcriptional misregulation in cancer of human) in the treatment of rhabdomyosarcoma. As shown in Fig 6, the activity of the hsa05202 pathway nodes were greatly inhibited for all eight cell lines of rhabdomyosarcoma, which demonstrated that the inhibition of hsa05202 pathway by drug CTK7H7014 has been successfully detected by pathDNN.

**Fig. 6:**
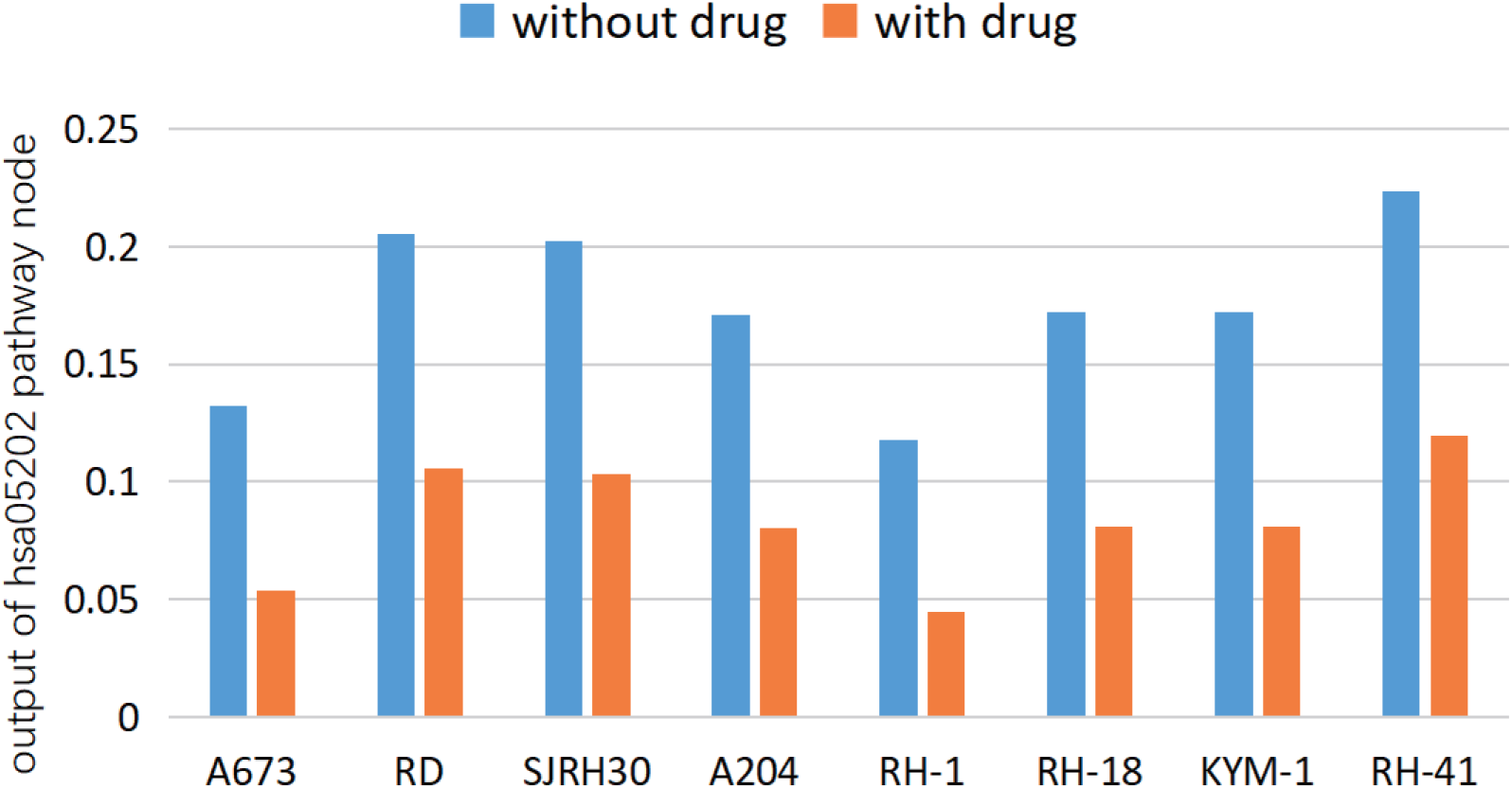
Significant decrease of the output of hsa05202 pathway node induced by CTK7H7014 on eight cell lines of rhabdomyosarcoma

**Fig. 7:**
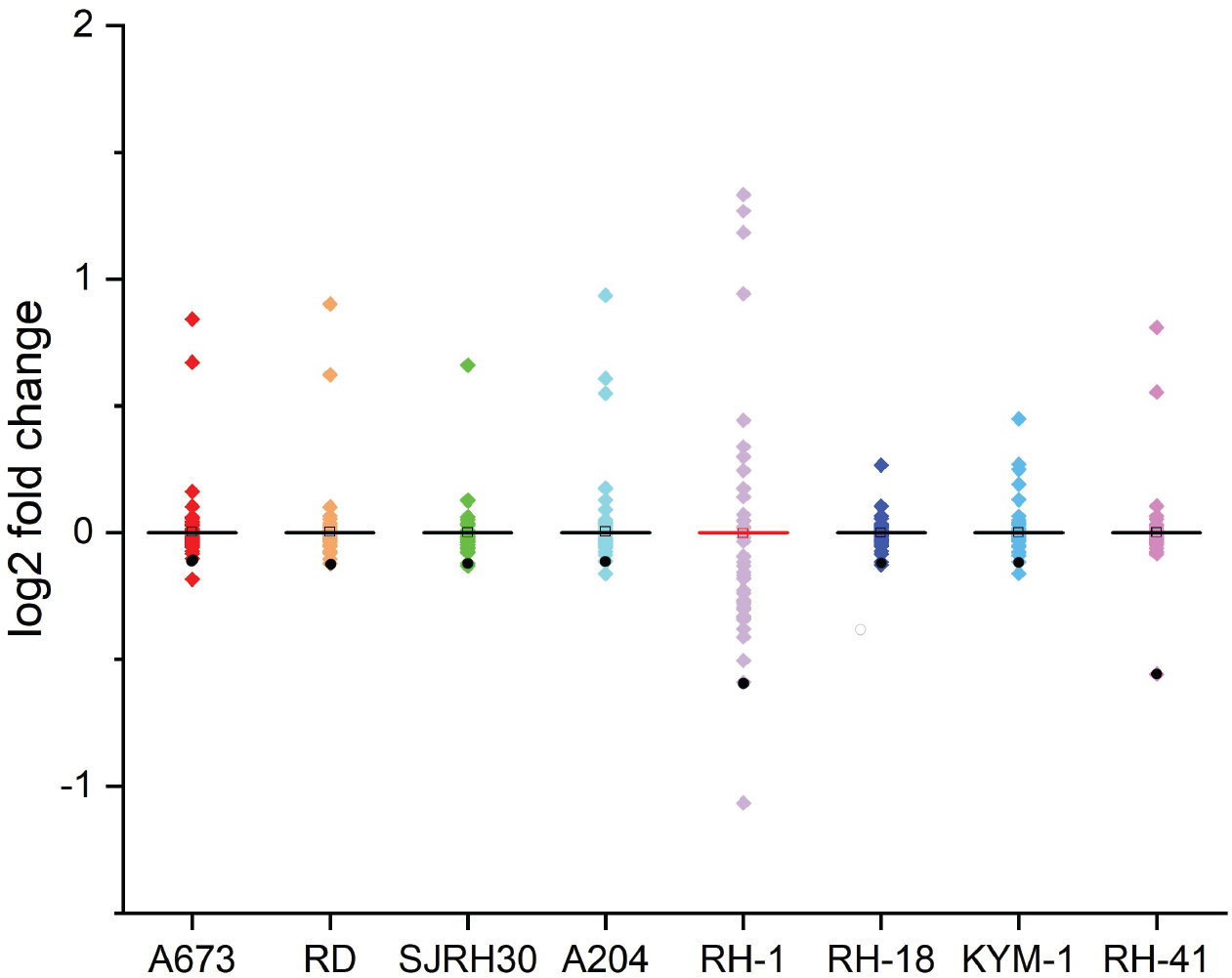
Boxplots of log2 fold changes of 323 pathway nodes induced by CTK7H7014 on eight cell lines of rhabdomyosarcoma

In addition, we also evaluated the activity change of all 323 pathway nodes in administration of drug CTK7H7014. To determine the level of change, we introduced log2 fold change that widely used in the expression analysis to represent the magnitude of change of pathway nodes. Intuitively, the smaller fold change is, the stronger the node is inhibited, and vice versa. As is illustrated in 7, the calculated log2 fold changes of all pathway nodes are presented in the box chart, the black point corresponds to the change of hsa05202 that is related to the disease rhabdomyosarcoma. Obviously, the inhibition of hsa05202 induced by CTK7H7014 is far more greatly inhibited compared to those of unrelated pathways, which demonstrates the excellent simulation and interpretability of our model.

### 3.7 Case studies

To demonstrate the practical application of pathDNN, we conducted two case studies from different perspectives. One aimed to find new drugs for certain cancer cell line, the other tried to find new indications for certain drugs. It is worth noting that the tested drug and cell line in the case studies were excluded from the training set. To discover new potential drugs for certain cancer cell lines, we exemplified T98G, a cell line of glioma that is a neurological disease. The top 10 drugs with the highest predicted sensitivity scores to T98G cell line by pathDNN are listed in Table 2. For each drug, we show its canonical name, predicted score and the remark that describes the evidence supporting its potential therapeutical treatment on glioma. Through retrieval of Pubmed, we found that 5 (Imatinib, Cyclopamine, Lapatinib, Erlotinib and Parthenolide) of these 10 drugs have solid literature support to treat glioma. For example, Ranza *et al.* reported that Imatinib in combination with gamma radiation has an additive antiproliferative effect on T98G [28]. Braun *et al.* assayed the transcriptionally active Gli1 in GBM cells and found that the presence of cyclopamine revealed a response to tumor cells [29]. Ramis *et al.* also found that EGFR inhibition Erlotinib can markedly inhibit cell motility and invasion in glioma cells [30]. In addition to the 5 aforementioned drugs, Crizotinib is currently under clinical trials for glioma registered in the clinicaltrais.gov database. Therefore, the predicted drugs are believed to be potential applications for glioma.

**Table 1:** Comparison of MSE loss and PCC between pathDNN and classical DNN

| model | MSE | PCC |
| --- | --- | --- |
| pathDNN | 0.014 | 0.8 |
| classical DNN (tuned hyperparameters and weights) | 0.018 | 0.73 |

**Table 2:** Top 10 drugs with the highest sensitivity scores for T98G cell line

| Drug | Predicted score | Evidence |
| --- | --- | --- |
| Crizotinib | 0.974 | under trial |
| Imatinib | 0.971 | pmid:20032406 |
| Cyclopamine | 0.97 | pmid:22406999 |
| Lapatinib | 0.969 | pmid:24815468 |
| GSK319347A | 0.967 |  |
| Erlotinib | 0.966 | pmid:22701710 |
| Parthenolide | 0.965 | pmid:29705182 |
| PHA-665752 | 0.965 |  |
| HG-5-88-01 | 0.964 |  |
| GNF-2 | 0.963 |  |

In another case study, we focused on the drug Etoposide, a semisynthetic derivative of podophyllotoxin that exhibits antitumor activity, which is widely used in leukaemias, lymphoma, lung cancer, ovarian cancer and other malignancies. To seek for its new indications, we listed the top 10 cancer cell lines with the highest predicted score by pathDNN in Table 3. Except for the names of cancer cell lines, we also presented associated disease names, predicted scores and supporting evidence. Trough retrieval of Drugbank, we found that NCI-H2066, NCI-H835, JHOS-4 and NCI-H250 are the cell lines derived from the cancers approved for administration by Etoposide. Besides, SW1116, CL-34 and CL-40, related to colon adenocarcinoma, and SK-MEL-31, related to melanoma, are currently under clinical trials to evaluate the therapeutical effect of Etoposide. Despite the lack of clinical support for the administration of gastric adenocarcinoma, Hotta *et al.* have demonstrated that Etoposide can effectively enhance the anti-tumor effects of Cisplatin in gastric cancer [31]. In addition, the anti-cancer activity of Doxorubicin and Etoposide on human breast cancer cells, ZR-75-30, has been verified to be enhanced with the participation of Polysaccharopeptide [32]. In conclusion, the predicted diseases or drugs significantly implicate the potential indications of new diseases, which are valuable for further clinical validation. The complete prediction results of T98G and Etoposide are presented in supplementary material s1.

**Table 3:** Top 10 cell lines with the highest sensitivity scores for Etoposide

| Cell line | Disease | Score | Evidence |
| --- | --- | --- | --- |
| SW1116 | Colon adenocarcinoma | 0.915 | under trial |
| KATOIII | Signet ring cell gastric adenocarcinoma | 0.892 | pmid:9137422 |
| SK-MEL-31 | Cutaneous melanoma | 0.877 | under trial |
| NCI-H2066 | Squamous cell lung carcinoma | 0.850 | pmid:28766886 |
| CL-34 | Colon adenocarcinoma | 0.834 | under trial |
| CL-40 | Colon adenocarcinoma | 0.833 | under trial |
| NCI-H835 | Lung carcinoid tumor | 0.824 | approved disease |
| JHOS-4 | High grade ovarian serous adenocarcinoma | 0.820 | approved disease |
| ZR-75-30 | Invasive ductal carcinoma | 0.819 | pmid:18292947 |
| NCI-H250 | Small cell lung carcinoma | 0.810 | approved disease |

## 4 Discussion and Conclusion

In this paper, we proposed a pathway-guided deep neural network model to predict drug sensitivity to cancer cells. To make the advantage of both the excellent predictive power of DNN and biological knowledge of pathways, we incorporated a layer of pathway nodes into the canonical DNN structure with custom connections between input gene nodes and pathway nodes, which makes the pathDNN model interpretable and predictive power. The extensive evaluations performed on several independent drug sensitivity datasets demonstrated that pathDNN significantly outperformed the canonical DNN model, as well as seven competitive methods, including three classical regression methods and four existing prediction methods. Most importantly, the remarkable output decrease of disease-related pathway nodes during forward propagation upon inputs of drug targets can imply the inhibition effect on corresponding pathways induced by drug treatment of cancer cells. In conclusion, the empirical experiments demonstrated that pathDNN achieves both pharmacological interpretability and predictive accuracy in modeling drug sensitivity to cancer cells.

By integration of our current knowledge presented in the form of pathways maps into the DNN structure, i.e. the pre-specified connections between the input layer and the pathway layer, pathDNN can extract the features in the pathway level, rather than gene level, to predict higher cellular process and final phenotypical changes via subsequent fully-connected layers. Our model actually functions similarly to the convolution neural network (CNN) widely used in image processing that extracts high features via a few lower convolutional blocks and then constructs higher representation of images. Accordingly, we observed remarkable activity changes of disease-related pathway nodes upon drug input signals, which implies significant impact on disease-related pathways induced by drug treatments with high sensitivity on cancer cells. In contrast, most of irrelevant pathway nodes show trivial changes in their activity. Our empirical experiments show that pathDNN achieves pharmacological interpretability and predictive ability in modeling drug sensitivity to cancer cells.

Last but not least, single-drug treatments often fail to inhibit the carcinogenic pathways in cancer cells, due to the intrinsic compensatory mechanism and cross-talk among cellular pathways. Combination drugs have demonstrated high sensitivities and low side effects in cancer therapies, and thus have drawn intensive attention from both the academical and industrial community [33, 34, 35]. It is crucial to develop *in-silico* methods that can dissect the mechanism of drug action from the perspective of pathways in modeling drug sensitivity to cancer. By virtue of the pharmacological interpretability, our pathDNN model can be easily extended to filter drug combinations that can cooperatively function to block carcinogenic pathways. In fact, we plan to develop a pathway-guided deep learning method to predict drug combinations in our near future works.

